# CD56 at the human NK cell lytic immunological synapse

**DOI:** 10.1101/2021.09.15.460290

**Authors:** Amera L. Martinez, Justin T. Gunesch, Emily M. Mace

## Abstract

CD56 is the main identifying cell surface molecule of NK cells and has been recently identified as a regulator of cytotoxic function in NK cell lines. Despite its newly defined role in lytic granule polarization and exocytosis, biological questions remain involving the localization and function of CD56 at the immunological synapse. Here we use confocal and structured illumination microscopy to demonstrate recruitment of CD56 to the peripheral supramolecular activating cluster (pSMAC) of the immunological synapse of lytic effector cells. We provide additional data demonstrating that cell lines that are less dependent on CD56 for function are not utilizing alternative pathways of cytotoxicity, and that those that are dependent on CD56 have normal expression of activating and adhesion receptors. Finally, we use actin reporter (LifeAct) expressing NK92 cell lines and live cell confocal microscopy to visualize live cell killing events with WT and CD56-KO cells. This work further characterizes the novel role for CD56 in cytotoxic function of NK cells and provides deeper insight into the role of CD56 at the NK cell immunological synapse.

## Introduction

Natural Killer (NK) cells are innate immune effector cells that are essential for the recognition and lysis of virally infected and tumorigenic cells. They are defined as CD56^+^ and CD3^−^ and make up 10% of lymphocytes circulating in peripheral blood. Through the formation of an immunological synapse with virally infected or tumorigenic cells, NK cells can mediate cytotoxic function, prompting the release of lytic granules that trigger target cell apoptosis. CD56 expression levels have been shown to correlate with differences in NK cell development, migration, and cytotoxicity (1, 2). However, the molecular function of CD56 in each of these contexts remains to be elucidated.

Target cell lysis by NK cells is dependent on the formation of the immunological synapse, which facilitates the polarization of activating receptors and the transport and release of perforin- and granzyme-containing lytic granules (3). The stages of immune synapse formation include cell adhesion mediated by LFA-1 and actin polarization, granule convergence to the microtubule organization center (MTOC), and MTOC polarization to the synapse (4). Calcium flux initiated by activating receptor ligation is necessary for the exocytosis of lytic granules, which occurs into the cleft between the NK cell and target cell (5, 6). Delivery of lytic granules or Fas-FasL or TRAIL aggregation can initiate target cell apoptosis (7, 8). Immune synapse formation includes the recruitment of multiple receptors for stabilization, activation and adhesion. One such receptor is CD56, which is enriched at the immune synapse of the NK92 cell line and required for its cytotoxic function (2).

CD56 (neural cell adhesion molecule, NCAM) is the major cell surface glycoprotein on human NK cells and on neuronal cells, where it facilitates adhesion (9, 10). Its extracellular Ig-like and fibronectin-binding domains mediate adhesion to the extracellular matrix through homophilic and heterophilic interactions (11, 12). In addition to its role in adhesion, CD56 can enable the NK cell-mediated lysis of CD56^+^ target cells (9, 12, 13). However, CD56 expression on target cells does not always increase NK cell cytotoxicity (14). In addition to the role of homophilic interactions with CD56 at the immune synapse, CD56 also serves as an NK cell pathogen recognition receptor against the CD56-negative fungi *Aspergillus fumigatus* where its recruitment to the synapse is actin-dependent (15). Additionally, CD56 facilitates the secretion of lytic granules and cytotoxic function against CD56-negative target cells by human NK cell lines through Pyk2 signaling (2).

To further describe the involvement of CD56 at the immunological synapse, we investigated requirements for activation, cytotoxicity, and recruitment of lytic granules after the deletion of CD56 in NK92 and YTS cell lines. Here, we provide additional insights into the differential requirement of CD56 for cytotoxic function of NK cell lines as described in Gunesch et al. 2020 (2) and the co-localization of CD56 and actin at the immunological synapse.

## Results

### Expression of CD18 and NKp30 on NK92 CD56-KO cell lines

Our previous study demonstrated that NK92 CD56-KO cells were impaired in their ability to mediate cytotoxicity despite intact conjugate formation with target cells (2). Cross-linking CD18 (LFA-1) and the activating receptor NKp30 leads to decreased secretion of lytic granule contents, suggesting that activation through these two receptors is impaired in CD56-KO cells, however we had not previously investigated the surface expression of CD18 and NKp30 on CD56-KO NK92 cells relative to WT NK92. We performed flow cytometry for CD18 and NKp30 and found that there was no significant difference in expression of either surface receptor between WT and CD56-KO NK92 cells (Fig. 1A). This result was consistent between technical replicates performed on different passages of cell lines (Fig. 1B). These data, in concert with our previous characterization of surface receptors on these cell lines (2), suggests that dysregulated activating receptor or integrin expression does not underlie the cytotoxicity defect found specifically in CD56-KO NK92 cells.

**Fig. 1.**
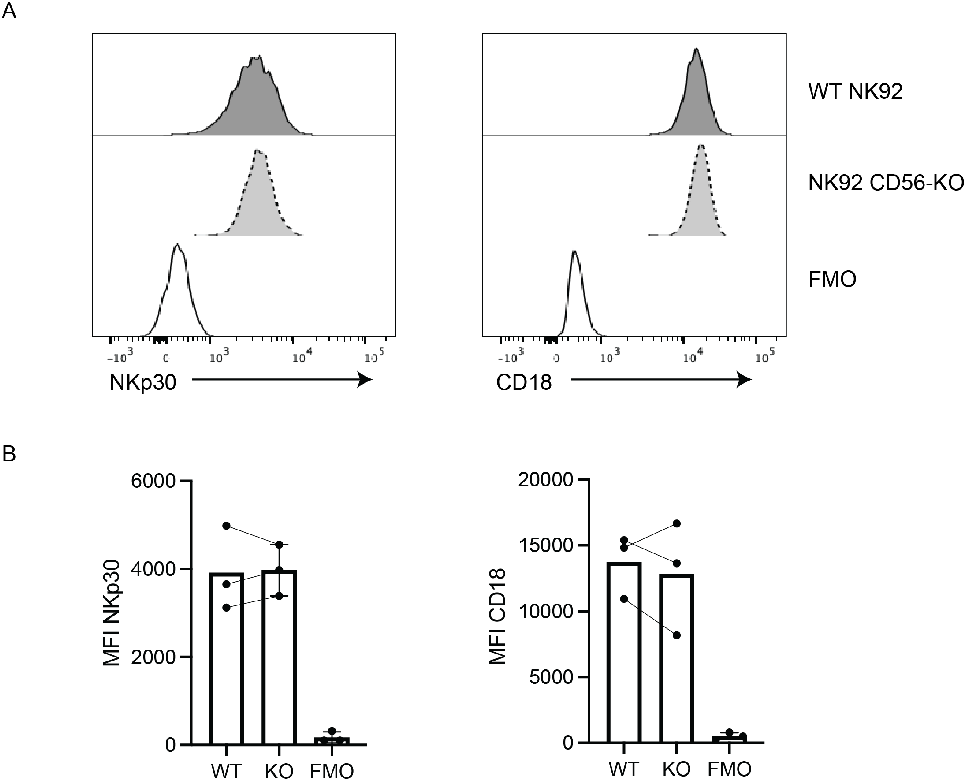
CD56 deletion does not affect the expression of CD18 or NKp30. WT or CD56-KO NK92 cells were immunostained for CD18 and NKp30 and analyzed by flow cytometry. A) Representative histograms from one experiment of 3 technical replicates. B) Mean fluorescence intensity (MFI) data from 3 technical replicates performed as independent experiments on different days. FMO, fluorescence minus one.

### Spatial localization of CD56 and actin at the NK cell lytic immune synapse

We previously have shown that CD56 is recruited to the immune synapse in NK92 and primary NK cells conjugated to susceptible target cells (2). To better define the spatial localization of CD56 at the immune synapse, we first performed additional analyses of the visualization of CD56 and actin from 3D volumetric images of NK cells conjugated to CD56-negative target cells. From images of WT NK92 cells conjugated to K562 targets we performed 3D reconstructions of the effector-target cell interface and rotated these to visualize the immune synapse ‘en face’ from the perspective of the target cell facing the effector cell synapse. We found that actin was enriched at the pSMAC of the immune synapse with perforin in the center (Fig. 2A). We also found that CD56 was enriched in this region and constructed a merged image to compare its localization to that of actin (Fig. 2B). A line profile drawn across the synapse provided intensity measurements for phalloidin (F-actin) and CD56 which confirmed this correlation (Fig. 2C). We then similarly conjugated primary ex vivo NK cells (eNK) to K562 targets to define localization of CD56. Despite their smaller size, eNK cells had consistent enrichment of CD56 at the peripheral supramolecular activating cluster (pSMAC) (Fig. 2D), which overlapped with actin localization (Fig. 2E). Intensity measurements also revealed similar peaks of actin and CD56 across the immune synapse as observed in NK cell lines (Fig. 2F). Lastly, we performed the same experiment with YTS cells conjugated to 721.221 targets. Despite their low reliance on CD56 for cytotoxicity (2), CD56 is found at the YTS cell immune synapse (Fig. 2G) where it localized at the pSMAC with actin (Fig. 2H). We further confirmed this localization via line profile measurements (Fig. 2I). These data suggest an association of CD56 with actin at the pSMAC of the lytic synapse that may be independent of a reliance on CD56 for target cell killing.

**Fig. 2.**
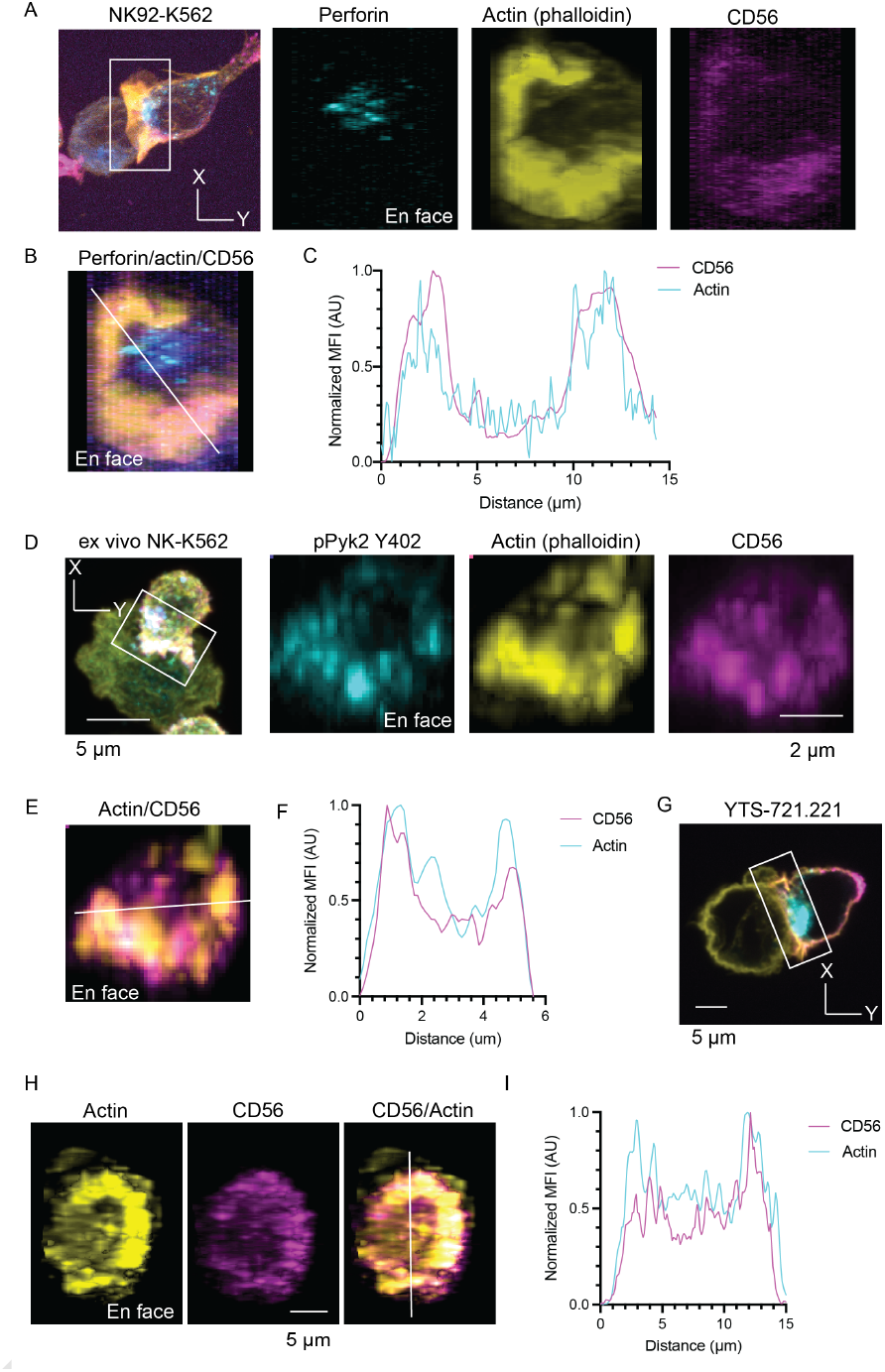
CD56 localizes to the pSMAC of immune synapses. WT NK92, ex vivo NK (eNK), and YTS cells were conjugated to K562 or 721.221 target cells. Conjugates were immunostained for perforin, actin, and CD56 and analyzed by confocal microscopy. A) Representative NK92:K562 conjugate shown in XY and YZ planes. B) Image at cross-section of synapse with a merge of all channels and diagonal line profile. C) Plot of mean fluorescence intensity of CD56 and actin across the line profile shown in (B). D) Representative eNK:K562 conjugate shown in XY and YZ planes. E) Image at cross section of synapse with a merge of all channels and diagonal line profile. F) Plot of mean fluorescence intensity of CD56 and actin across the line profile shown in (E). G) Representative YTS:721.221 conjugate immunostained for perforin (cyan, actin (yellow) and CD56 (magenta) and shown in XY plane. H) Image at cross-section of synapse with a merge of CD56 and actin and line profile marked. I) Plot of mean fluorescence intensity data of CD56 and actin across the line profile shown in (H). Data representative of 3 independent experiments and 3 biological donors (eNK cells).

### High-resolution imaging confirms decreased actin-CD56 colocalization in YTS cells

To better define the localization of CD56 in response to activating signaling, we activated NK cells on functionalized glass surfaces to recapitulate the immune synapse and imaged cells using structured illumination microscopy (SIM). NK92 cells spread symmetrically on anti-CD18/NKp30 coated surfaces and remodelled actin to form a lamellipodia and a cortical actin meshwork that is permissive for degranulation (Fig. 3A) (16–18). As suggested by our fixed cell conjugates imaged by confocal, we found CD56 strongly associated with actin, particularly in regions of actin density at the outer edge of the synapse. We further imaged CD56 and actin at the plane of the glass by SIM in YTS cells following activation by CD18 and CD28 cross-linking. As in NK92 cells, CD56 was found particularly localized to cortical actin and along filopodia protruding from the synapse (Fig. 3B). Line profiles drawn across the synapse of both NK92 and YTS cells confirmed that sites of actin enrichment were frequently also enriched for CD56 in both cell lines; however, this localization was not as strong in YTS cells as it was in NK92 cells (Fig. 3C). These data demonstrate that the colocalization of actin and CD56 that we observed in the periphery of the immune synapse of NK cells conjugated to target cells could be recapitulated using activation on functionalized glass and further showed that this apparent colocalization of CD56 and actin was detectable by enhanced resolution microscopy.

**Fig. 3.**
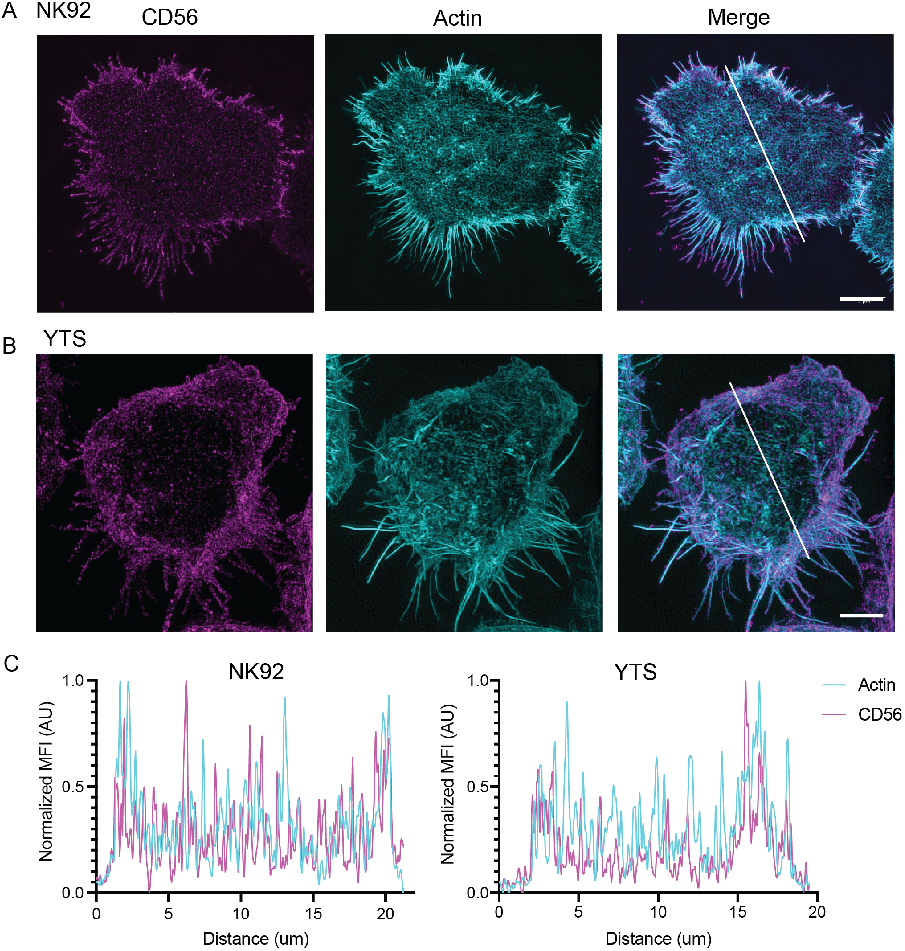
CD56 colocalizes with actin at the immune synapse. WT cells were immunostained for CD56 and actin and imaged by structured illumination microscopy. A) Representative images of WT NK92 cells at the plane of activated glass. Representative images of WT YTS cells at the plane of activated glass. C) Plot of mean fluorescence intensity data taken from line profiles of NK92 (left) and YTS (right) cells shown above. Scale bar = 5 μm. Images are maximum intensity projections of 3 glass-proximal 0.025 μm Z slices.

### Requirement for calcium flux in cytotoxic function of YTS CD56-KO cell lines

Having identified CD56 at the immune synapse in both NK92 and YTS cells, we sought to better understand the differential requirement for CD56 in NK cell-mediated target cell lysis. Our group previously identified a crucial role for CD56 in lytic granule exocytosis and cytotoxicity of NK92 cells, whereas YTS cells had impaired IFNγ secretion yet retained some lytic function following deletion of CD56 (2). While the primary mechanism for NK cell cytotoxicity is thought to be lytic granule exocytosis, alternative forms of target cell killing include Fas-FasL and TRAIL-mediated receptor-ligand interactions (7, 8). To determine whether YTS cells were compensating for loss of CD56 expression by mediating target cell killing through these alternative pathways, we used the calcium chelator BAPTA-AM to test the requirement for calcium flux in the context of YTS cytotoxicity. We performed a four hour chromium release assay in the presence of this calcium chelator. We found that YTS cells had a requirement for calcium flux in exocytosis, as WT YTS had decreased cytotoxicity in the presence of 20μM BAPTA-AM and increasing the concentration of BAPTA-AM to 40μM produced an almost absolute defect in cytotoxicity (Fig. 4A). We also found a similar impairment of target lysis with calcium chelation of CD56-KO YTS cells, which were previously shown to have decreased, but not absent, cytotoxic function (2) (Fig. 4A). These data demonstrate that both cell lines rely on calciumdependent signaling signaling to lyse targets and expand our understanding of the effect of CD56 deletion on YTS cells. Therefore, the residual cytotoxic function of CD56-KO YTS cells is not a result of a calcium-independent pathway of exocytosis in YTS cells. CD56-KO YTS cells have decreased cytotoxic function after a 1 hour incubation with 721.221 target cells (2), but we wanted to additionally test the reliance of YTS cells on CD56 for cytotoxicity at different effector to target cell ratios. Notably, we found that CD56-KO YTS cells had decreased cytotoxic function when compared to WT YTS cells when we seeded the assay at a lower (5:1) effector to target cell ratio (Fig. 4B), however cytotoxicity was still not profoundly decreased as it consistently is with NK92 CD56-KO NK cells (2).

**Fig. 4.**
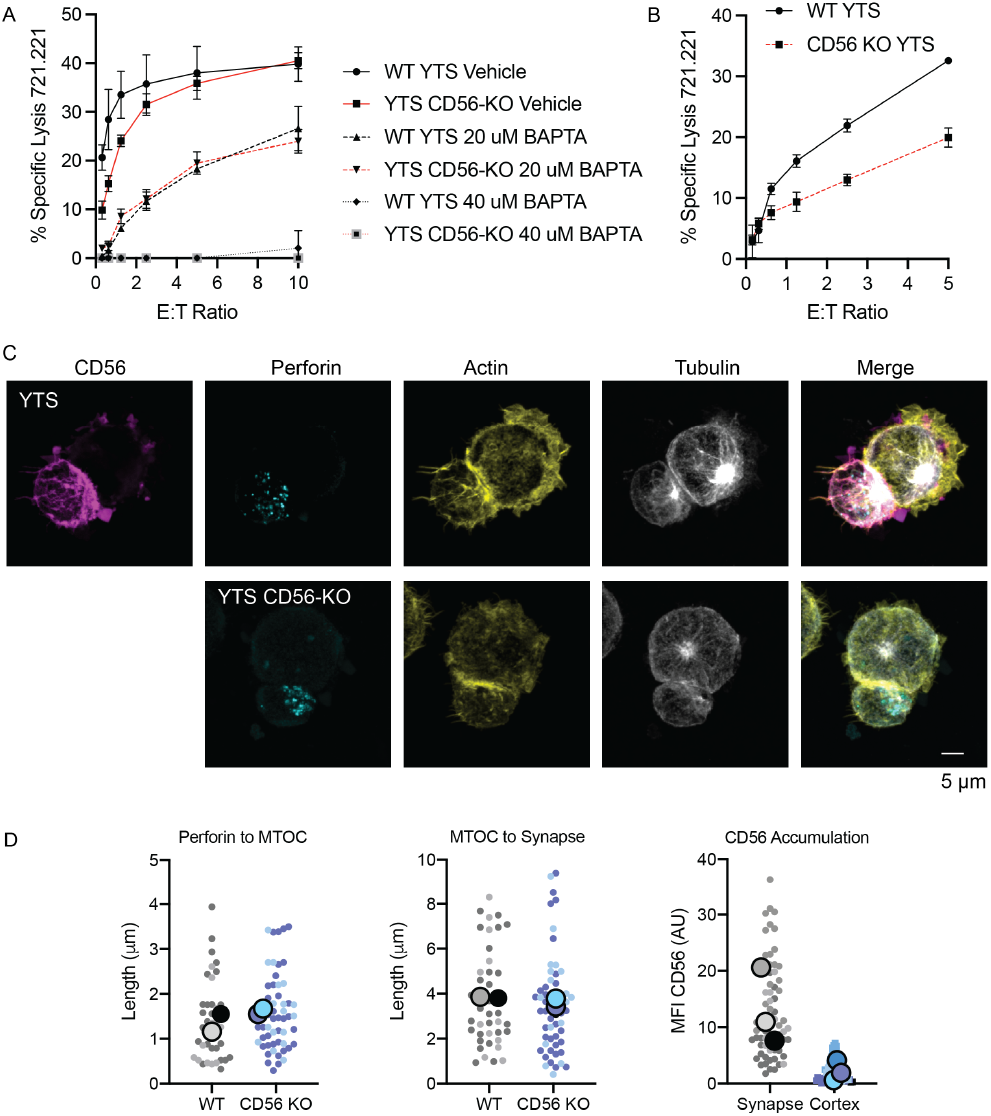
CD56 is enriched at the synapse of YTS cells which lyse targets in a calcium-dependent manner. WT and CD56-KO YTS cells were conjugated to 721.221 target cells and analyzed by 51Cr-release assay and microscopy. A) 4 hr 51Cr assay performed with WT and CD56-KO YTS cells with vehicle control (DMSO) or 20 or 40 μM of cell-permeable calcium chelator, BAPTA-AM, as effectors against 721.221 targets. Data shown are the mean of triplicates from one representative of 3 independent experiments. B) 4 hr 51Cr assay performed as in (A) but with a 5:1 effector to target cell ratio. C) Representative images from >30 cells taken by confocal microscopy of WT (top) and CD56 KO (bottom) YTS cells conjugated to 721.221 targets in a 2:1 ratio for 45 min before fixation, immunostaining as indicated, and imaging. C) Image quantification for distance of perforin centroid to MTOC (left), distance of MTOC to immune synapse (middle), and integrated intensity of CD56 at the immune synapse (right). Data are displayed as a SuperPlot where large dots symbolize the mean of three technical replicates for CD56 accumulation, n=67; and two technical replicates for perforin to MTOC and MTOC to synapse measurements, n = 39 (WT), n = 57 (KO).

Having confirmed that YTS cells execute cytotoxicity through calcium-dependent lytic granule exocytosis and given the impaired synapse formation of NK92 cells (2), we sought to determine whether immune synapse formation was impaired in CD56-KO YTS cells. Confocal microscopy of YTS cells conjugated to 721.221 target cells allowed for visual comparison of CD56-KO YTS cells to their WT counterparts (Fig. 4C) which displayed normal perforin convergence and polarization towards the immune synapse. In addition, MTOC polarization and actin accumulation appeared to be normal in CD56-KO YTS cells. These images were then quantified for perforin convergence, MTOC polarization, and CD56 accumulation at the immune synapse (Fig. 4D). These analyses demonstrated no significant change in the distance of perforin-containing lytic granules to the MTOC or distance in MTOC to the immune synapse in CD56 KO YTS cells. Consistent with our observation that actin and CD56 colocalized to the YTS lytic immune synapse (Fig. 2), we also found that WT YTS cells had significantly increased CD56 intensity at the synapse of conjugated cells when compared to the distal cortex.

### Requirement for CD56 in granule polarization in NK92 cells

The deletion of CD56 in NK92 cell lines was previously shown to contribute to impaired granule polarization towards target cells (2). We wanted to model this in a cell-free manner by imaging WT and CD56-KO NK92 and YTS cells in three dimensions on functionalized glass and confirm that our activating conditions result in granule polarization to the glass in each cell line. Antibodies to activating receptors CD28 (YTS) or NKp30 (NK92) and the integrin LFA-1 (CD18) were coated on glass and cells were stained for actin and perforin after spreading for 20 minutes (Fig. 5A). Cells were imaged with 3D confocal microscopy starting from the plane of the activated glass. After visualization in Z and Y, WT NK92 and YTS as well as CD56-KO YTS all had clear convergence and polarization of lytic granules towards the glass. However, NK92 CD56-KO cells lacked granule convergence and polarization, with lytic granules distributed throughout the cytoplasm (Fig. 5A). By bisecting each volumetric cell image, we quantified the density of perforin in either the glass-proximal, which we refer to as synapse, or glass-distal, cell sections (Fig. 5B, C). Intensity measurements confirmed a greater amount of perforin in the glass-distal region of CD56-KO NK92 cells than in other cell types. The intact polarization of CD56-KO YTS cells when compared to CD56-KO NK92 cells further demonstrates that formation of the YTS cell immune synapse is not CD56-dependent. Further, the polarization of NK cells under these conditions formally demonstrates that the secretion induced by these conditions is directional and effectively models the NK cell lytic immune synapse.

**Fig. 5.**
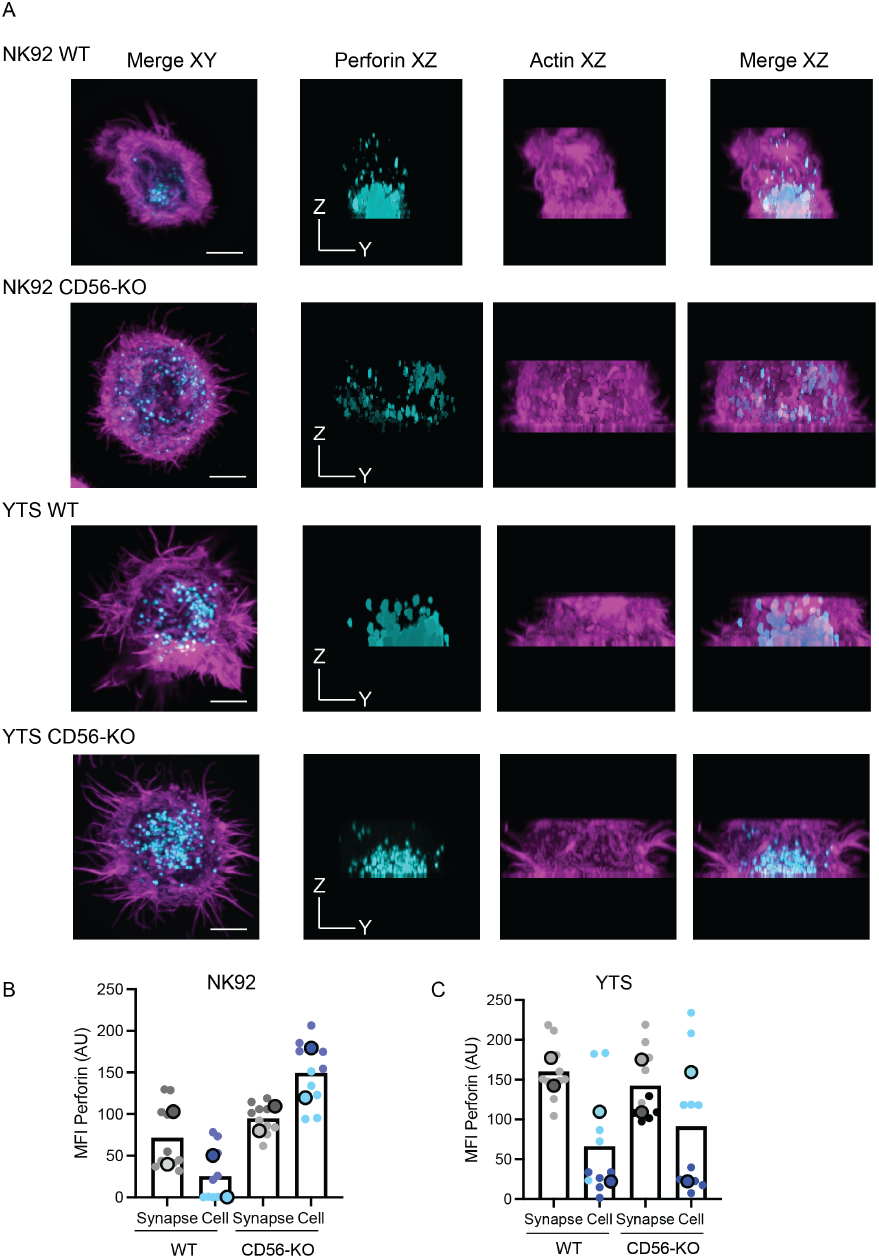
CD56-KO NK92 cells have impaired granule convergence and polarization. WT or CD56-KO NK92 and YTS cells were immunostained for perforin and actin and analyzed by fixed cell confocal microscopy. A) Representative images from one experiment from 2 technical replicates shown at the plane of glass coated with anti-CD18 and either anti-NKp30 (NK92) or anti-CD28 (YTS) in XY (left) and YZ orientation. B) Mean fluorescence intensity data of perforin from 2 technical replicates (n = 12) compared between the synapse and cell of WT and CD56-KO NK92 cells. C) Mean fluorescence intensity data of perforin from 2 technical replicates (n = 12) compared between the synapse and cell of WT and CD56-KO YTS cells. Scale bar = 5 μm.

### Delayed target cell lysis by CD56-KO NK92 cells

Our previous studies of NK cell line conjugation and cytotoxicity showed defects in the function of NK92 CD56-KO cells (2). Namely, target cell lysis, but not conjugation, was impaired at 1 and 4 hour time points and linked to impaired lytic granule exocytosis. However, these assays performed at set time points could miss certain kinetics of cytotoxic function. Therefore, we sought to visualize cytotoxicity of WT and CD56-KO NK92 LifeAct mScarlet cell lines in real time using confocal microscopy. Live cell killing assays were performed by labeling LifeAct mScarlet expressing NK92 effector cells with a pH sensitive indicator of lytic granule lysosomes (Lysotracker Deep Red) and target cells with a cytoplasmic dye that is lost upon disruption of the cell membrane, marking cell death (Calcein Green). Imaging of WT and CD56-KO NK92 conjugated to K562 cells at two minute intervals enabled us to identify the formation of conjugates and define the remodeling of actin at the immune synapse, the polarization of lytic granules, and target cell death. These experiments consistently demonstrated a lack of cell death at 30 minutes in CD56-KO cells, whereas WT cells effectively killed their target (Fig. 6A, Supplemental Videos 1 and 2). However, we frequently visualized CD56-KO cells in close proximity or conjugated to target cells while failing to undergo extensive actin remodeling or granule polarization seen in WT cells. While in some cases we did observe target cell lysis by CD56-KO cells, it was delayed relative to WT cells. These data further validate the findings from our fixed cell microscopy and add additional depth to our understanding of the dynamics of immune synapse formation in the absence of CD56 expression.

**Fig. 6.**
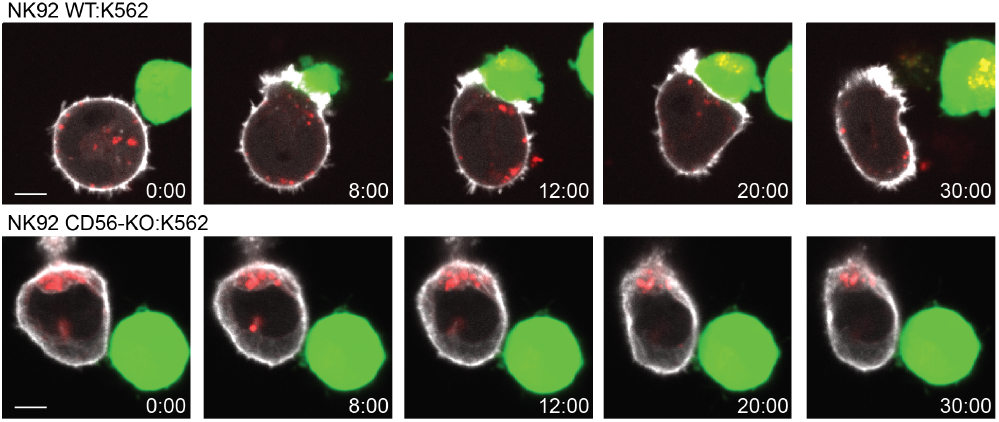
CD56-KO NK92 cells have a delayed ability to lyse target cells. WT or CD56-KO NK92 cells expressing LifeAct mScarlet to visualize actin were co-cultured with K562 target cells and analyzed by live cell confocal microscopy. A) Representative images isolated from time series of one experiment from 3 technical replicates showing WT (top) and CD56-KO (bottom) NK92 LifeAct mScarlet (grey) effectors stained with Lysotracker Deep Red (red) conjugated to K562 targets stained with Calcein green (green). Scale bar = 5 μm. See also Supplemental Videos 1 and 2.

## Discussion

Despite being the commonly used identifier of human NK cells, the functional roles of CD56 are still being uncovered. Previous studies showed a requirement for homotypic interactions between CD56 on the NK cell and target cell for cytotoxicity (9), whereas another group found that this was not required as they saw no difference in the lysis of CD56+ and CD56 target cells (14). Treatment with monoclonal antibodies against CD56 diminishes target cell recognition and cytotoxicity by NK cell lines (19) and CD56 expression in a breast cancer model increases cytotoxicity of NK92 cells (13). However, in some contexts, NCAM expression on target cells can promote tumor cell survival, in part through the prevention of conjugate formation with NK cells (12). While these studies suggested that CD56-CD56 interactions may play a role in mediating cytotoxicity under certain conditions, the deletion of CD56 in human NK cell lines has been shown to eliminate or delay target cell lysis, impair immunological synapse formation, and decrease secretion independently of NCAM-NCAM interactions (2). As the cytolytic utility of NK cells shows promise as a cancer immunotherapy, further defining the functional role of CD56 at the immunological synapse will both increase our understanding of basic NK cell biology and the therapeutic applications of human NK cells..

Our previous studies have defined a requirement for CD56 in NK92 cell migration and cytotoxicity, in part by analyzing the effect of CRISPR-Cas9 deletion of CD56 in NK92 and YTS cell lines (1, 2). Here, we aimed to expand upon the function of CD56 in NK cell cytotoxicity by analyzing the role that it plays at the NK cell lytic immune synapse. In addition, we sought to better understand the differential requirement for CD56 in YTS and NK92 cytotoxic function. Our experimental model includes the activation of NK cell lines using cross-linking with activating antibodies on coverslips, which has been previously shown to recapitulate immune synapse formation and NK cell degranulation (17). However, despite no apparent dysregulation of the NK cell surface receptor phenotype in CD56-KO cells, we had not previously validated expression of the receptors utilized in our NK92 activation model, namely CD18 and NKp30. Given the importance of LFA-1 in NK cell immune synapse formation and function (3), we sought to confirm that any immune synapse defects present in CD56-KO cell lines were not the result of loss of LFA-1 expression. The normal LFA-1 expression on CD56-KO NK92 cells is consistent with our previous finding that CD56-KO NK92 cells form conjugates with K562 targets at equal or higher frequency than WT cells at 30 and 60 minutes (2). Collectively, our studies suggest that LFA-1-mediated adhesion is intact in CD56-KO cells and further suggests that CD56 does not mediate adhesion to target cells. Therefore, while NCAM can mediate cell adhesion through homophilic and heterophilic interactions (11, 12), it does not appear to play a role in NK cell adhesion either directly or by affecting integrin expression. However, the expression of CD56 on NK cell exosomes and the requirement of CD56 for exosome adhesion to target cells (20, 21) emphasizes the need for further studies to fully understand the role of CD56 in immune cell interactions.

To identify differences that could elucidate requirements of CD56 for cytotoxic function, we directly compared the NK92 and YTS cell lines through the imaging of the immunological synapse. En face projections of WT NK92, primary NK cells, and YTS cells conjugated to target cells showed consistent localization of perforin, actin, and CD56 at the immunological synapse. Interestingly, CD56 appeared to be associated with actin at the pSMAC of the immunological synapse. Previous studies have demonstrated an association of CD56 with actin while serving a role as a pathogen recognition receptor. After treatment with an actin polymerization inhibitor, human NK cells demonstrated reduced CD56 recruitment to the site of contact with *Aspergillus fumigatus* (15), suggesting that actin remodeling is required for CD56 localization. Our analysis done in a target cell-free model with activating antibody-coated glass and super resolution microscopy confirmed the association between actin and CD56 in both NK92 and YTS cells. However, there appeared to be less correlation between the localization of CD56 and actin in YTS cells than in NK92 or primary NK cells. While it is unclear what the link is between CD56 and actin, NCAM is required for the recruitment and activation of focal adhesion kinase (FAK) for neurite outgrowth of neuronal cells (22). Pyk2 is a FAK homologue that is expressed in NK cells and regulates Rac activation required for actin polymerization (23–25). Pyk2 also mediates the regulation of cytotoxicity by CD56 in NK92 cells, which have a higher dependence on Pyk2 function than YTS cells (2). The observed decrease in association of CD56 and actin in YTS cells may therefore reflect a differential usage of Pyk2 by these two cell lines.

Our previous study showed a decrease in cytotoxicity after 4 hours of NK92 CD56-KO cells with target cells but YTS CD56-KO cells only had decreased killing at a 1 hour time point (2). We speculated that the reduced requirement of CD56 for cytotoxic function could be attributed to YTS cells utilizing alternative pathways for cytotoxicity from NK92 cell lines. This led us to investigate a differential requirement for calcium between the two NK cell lines for target cell lysis. 51Cr release assays performed on WT and CD56-KO YTS cells that were untreated or treated with a cell-permeable calcium chelator demonstrated that YTS cells lyse 721.221 targets in a calcium-dependent manner. Additional 51Cr release assays performed on the same cells at a lower effector cell density showed a decrease in cytotoxic function of CD56-KO YTS cells. This implies that the mechanism in which YTS cells are able to lyse target cells in the absence of CD56 is less compensatory when there are fewer effector cells. This observation, taken together with the impaired lysis of target cells by YTS CD56-KO cells at one hour, but not four hour timepoints, may be linked to dynamics of target cell killing including migration or detachment, especially given the role that CD56 plays in NK cell migration (1). Together with the impairment of granule polarization in NK92 CD56-KO cells, but not YTS CD56-KO, this suggests that the greater ability of YTS CD56-KO cells to kill targets relative to NK92 CD56-KO cells is related to differences in activation and exocytosis between the cell lines and not the utilization of alternative forms of killing by YTS cells. Despite this intact activation, however, there remain differences in the dynamics of killing by the YTS CD56-KO cells that are evident in contexts of shorter incubation times or lower cell densities.

Fixed cell confocal microscopy of YTS and primary NK cell effectors conjugated to 721.221 targets showed an enrichment of CD56 at the immunological synapse, and it is interesting that CD56 is recruited to the synapse of NK cells that do not appear to rely on it as extensively for cytotoxicity. Confocal microscopy images also demonstrated no impact of CD56 deletion on the convergence and polarization of perforin and the MTOC to the YTS immune synapse. Our previous work in NK92 conjugates also showed no effect on granule convergence but there was a significant defect in granule polarization with CD56 deletion, which we further confirmed here using confocal imaging of cells on functionalized glass. This points to a requirement of CD56 for the movement of the MTOC and associated lytic granules to the synapse in NK92, but not YTS, cells. Lytic granules can converge independently of MTOC polarization and actin reorganization and in events that do not lead to target cell death, further demonstrating that early signals leading to convergence, such as LFA-1 mediated signaling leading to cell adhesion and granule convergence, are not impaired in CD56-KO NK92 cells (26, 27). The dependence of lytic granule convergence but not MTOC polarization and actin reorganization on Src-kinase signaling suggests another avenue for CD56 function (28). The requirement of CD56 for MTOC translocation in NK92 cells aligns with the recruitment of Pyk2 to the immune synapse that is required for the MTOC polarization necessary for cytotoxic function (29). Given the greater dependence of NK92 cells on Pyk2 for cytotoxicity (2), these findings elucidate a potential pathway for the impaired MTOC polarization in CD56 KO-NK92 cells that could operate in a CD56-independent manner in YTS cell lines.

Our target cell-free model enabled us to more closely examine the localization of perforin in WT and CD56-KO NK92 and YTS cells via confocal microscopy and identified a lack of polarization of perforin-containing granules only in CD56-KO NK92 cells in response to cross-linking of integrin and activating receptors. While there were granules near the synapse of CD56-KO NK92 cells, convergence and polarization was disrupted compared to the other cell types. It is important to note that degranulation of a single perforin-containing granule can lead to target cell death (30), so the observation of some synaptic granules, even in CD56-KO NK92 cells, could explain why there is not a total loss of cytotoxic function in CD56-KO cell lines. Alternatively, CD56 may be playing another novel role in lytic granule dynamics. Previous studies defining roles for NCAM in exocytosis and vesicle trafficking underscore the numerous roles it can play in functions that could be relevant to NK cell biology.

Lastly, it can be difficult to fully understand defects in the dynamic processes of cytotoxicity with fixed cell microscopy. To better understand the kinetics of cytotoxicity in WT and CD56-KO NK cell lines, we performed a live cell killing assay imaged via confocal microscopy using cells expressing a LifeAct reporter to visualize actin and labeled with Lysotracker to monitor lytic granule movement. As expected, NK92 cells consistently killed target cells within 20-25 minutes, whereas CD56-KO NK92 cells failed to lyse targets. Similarly to our fixed cell microscopy data, granules failed to polarize toward target cells despite conjugates being formed. CD56-KO NK92 cells often made contact with a target, yet rolled around with it without lysing the target cell. This observation aligns with our previous study which demonstrated an impact on the cytotoxic effector function of CD56-KO cell lines after 1 hour (2). We observed a few instances of target cell lysis by CD56-KO NK92, however in these cases the effector failed to release its target. This suggests that CD56 deletion causes prolonged synapse time, which has been shown to cause hypersecretion of inflammatory cytokines which can be detrimental to human health (31, 32). The failure to detach from target cells could also interfere with the known serial killing function of NK cells which contributes greatly to their use in cell-based therapies for tumor reduction (33). Another interesting observation of CD56-KO NK92 cells was the lack of lamellipodial ruffling at the immunological synapse that was visualized in the WT NK92 cells, signifying a lack of commitment to a full synapse. Our assay involved the plating of effectors with targets in a 2:1 ratio, however, it would be very useful to use this model with a microwell to separate the effector and target by a distance large enough to require that effectors migrate to reach their target. Since CD56 is required for NK cell migration (1), this would be important in defining the contribution of a migratory defect to the cytotoxicity defect. Taken together, our data demonstrate that NK92 cells recruit CD56 to the immunological synapse and play a role in target cell lysis which seems to intertwine with actin dynamics at the synapse.

In summary, here we further describe the role of CD56 in cytotoxicity and the differential requirements for this molecule between two commonly used NK cell lines. YTS cells kill in a calcium-dependent manner and have enriched CD56 at the immunological synapse. As these cells have less dependence on CD56 for cytotoxic function, this enrichment suggests that CD56 may be recruited to the immune synapse independently of its function there, which is also supported by our data showing that it is recruited to the synapse of ex vivo NK cells, which also do not require CD56 for function (2). YTS cells do not demonstrate defects in lytic granule polarization with CD56 deletion, whereas NK92 cells do, again suggesting less of a reliance on CD56 for signaling in YTS cells despite the decreased function of CD56-KO YTS cells measured by Cr51 assay. These findings lend additional insight into the complex functions of CD56.

## Methods

### Cell lines and cell culture

Cell lines were cultured at 37°C 5% CO2 at an approximate concentration of 10^5^ cells/ml. YTS, 721.221 and K562 cells were a gift from Dr. Jordan Orange (Columbia University) and NK92 cells were acquired from the ATCC. YTS cells were cultured in RPMI media supplemented with 10% fetal bovine serum and NK92 cells were cultured in Myelocult (Stemcell Technologies, catalogue H5100) supplemented with 200 U/ml of IL-2 (Roche) and 50 U/ml of penicillin/streptomycin (Gibco). Cell line validation was routinely performed by flow cytometry for cell surface receptors, with NK92 cells defined as being CD56^+^CD3^−^CD2+ and YTS cells as CD56^+^CD3^−^CD2-. NCAM1 (CD56) was deleted from NK92 and YTS cells by CRISPR-Cas9 as previously described (2). LifeAct mScarlet cell lines were generated by electroporating 2 μg/ml of pLifeAct-mScarlet-N1 plasmid DNA (a gift from Dorus Gadella, Addgene plasmid 85054) into WT and CD56-KO NK92 cells using the Amaxa nucleofector (Kit R) then growing in the presence of 1.3 mg/ml of Geneticin (G418, Fisher Scientific) to select for transfected cells and FACS sorting for fluorescent signal. CD56-KO cells were frequently validated by flow cytometry for the absence of CD56 and presence of other defining NK cell surface markers and all cell lines were routinely confirmed to be mycoplasma negative.

### Flow cytometry

Flow cytometry was performed on cell lines as independent technical replicates with different passages of cells on different days. NK92 (WT or CD56-KO) cells were harvested, washed once with complete media and resuspended in PBS for immunostaining. Cells were incubated with Brilliant Violet 421 anti-CD56 (clone HCD56, Biolegend, 3 μg/ml), APC anti-NKp30 (clone P30-15, Biolegend, 10 μg/ml) and FITC anti-CD18 (clone 6.7, BD Pharmingen, 1:100) for 30 minutes at 4 deg C in the dark then washed once with PBS. Fluorescence minus one controls were prepared for NKp30 and CD18 conditions. Data were acquired on a BD Fortessa cytometer and exported to FlowJo (BD Biosciences) for analysis and graphed using Prism 8.0 (GraphPad).

### Chromium Release Assay

721.221 target cell lines were incubated with 100 μCi 51Cr for 1 hr at 37°C and 5% CO2. These cells were then co-cultured with YTS effector cells in 96-well round-bottom plates for 4 hrs at 37°C and 5% CO2. Following incubation, total release control target cells were lysed with 1% IGEPAL (Sigma Aldrich) and plates were centrifuged, then supernatant was harvested and transferred to LUMA plates (Perkin Elmer). After drying overnight, plates were read with a TopCount NXT. For analysis, % specific lysis was calculated by: (sample - avg spontaneous release) / (avg total release - avg spontaneous release) * 100. For experiments done with calcium chelation, assays were done in the presence of 20 μM or 40 μM BAPTA AM (ThermoFisher) or vehicle control (DMSO). All assays were performed in triplicate, with independent replicates performed on different days with different passages of cell lines.

### Confocal microscopy

For conjugate assays, WT NK92, eNK, and WT and CD56-KO YTS cell lines were co-cultured with K562 (NK92, eNK) or 721.221 (WT and CD56-KO YTS) target cells at a 2:1 effector to target ratio in complete RPMI 10% FBS media. Cells were incubated for 20 mins at 37°C 5% CO2 in 1.5ml microcentrifuge tubes. Conjugates were then gently mixed and transferred to 0.01% poly-L-lysine coated #1.5 coverslips. Coverslips were incubated for 25 mins at 37°C 5% CO2 and then fixed and permeabilized with CytoFix/CytoPerm (BD Biosciences) at room temperature for 20 mins. Fixative was removed and coverslips were rinsed three times with 150 μl PBS. Immunostaining was performed with biotinylated monoclonal mouse antitubulin (clone 236-10501, Thermo, 1 μg/ml) then Brilliant Violet 421-conjugated streptavidin (Biolegend, 0.1 μg/ml), Alexa Fluor 488-conjugated mouse anti-perforin (Biolegend, clone dG9), and phalloidin Alexa Fluor 568 (Thermo). Prior to conjugation, NK cells were pre-incubated for 20 minutes with anti-CD56 Alexa Fluor 647 (HCD56, Biolegend, 5 μg/ml). For granule polarization assays, WT and CD56 KO NK92 and YTS cells were incubated for 20 mins at 37°C and 5% CO2 on .5 glass coverslips that had been pre-coated with 10 μg/ml anti-CD18 (clone IB4) and anti-NKp30 (clone P30-15, Biolegend, NK92) or anti-CD28 (clone CD28.2, BD Biosciences, YTS). NK cells were fixed and permeabilized with CytoFix/CytoPerm (BD Biosciences) at room temperature for 20 mins. Fixative was removed and coverslips were rinsed three times with 150 μl PBS. Immunostaining was performed with Alexa Fluor 488-conjugated mouse anti-perforin (Biolegend, clone dG9) and phalloidin Alexa Fluor 568. Coverslips were mounted on slides with ProLong Glass antifade reagent (ThermoFisher Scientific). Images were acquired with a 63X 1.40 NA or 100X 1.46 NA objective on a Zeiss AxioObserver Z1 microscope stand equipped with a Yokogawa W1 spinning disk. Illumination was by solid state laser and detection by Prime 95B sCMOS camera. Data were acquired in 3i Slidebook software and exported as TIFF files for further analysis.

For live cell killing assay, WT and CD56-KO NK92 cells expressing LifeAct mScarlet were incubated with LysoTracker DeepRed and K562 cells were incubated with Calcein Green for 1 hr at 37°C and 5% CO2 in 15 ml conical tubes with loose caps. 8 well chamber slides were coated with 0.01% poly-L-lysine for 1 hr at room temperature then washed vigorously with deionized water and PBS three times each. Chamber slides were then coated with 5 μg/ml fibronectin for 30 mins at 37°C and 5% CO2. Cells were washed in media and added to the wells of chamber slides at a 3:1 effector to target ratio. Images were acquired with a 63X 1.40 NA confocal objective using the 5X5 montage setting of the 3i Slidebook software. A range of 0.17μm Z step sizes were imaged at an interval of 2 minutes for 1 hour. The 647 and 488 lasers were used and set to 3% power with the lowest exposure possible along with imaging in brightfield. Data was exported as .tiff files (XYZT) for further analysis.

### Structured illumination microscopy

NK92 and YTS cell lines were incubated for 20 mins at 37°C and 5% CO2 on #1.5 glass coverslips that had been pre-coated with anti-CD18 (clone IB4, 10 μg/ml) and 10 μg/ml anti-NKp30 (clone P30-15, Biolegend, NK92) or anti-CD28 (BD Biosciences, YTS). NK cells were pre-incubated with CD56 Alexa Fluor 488 (HCD56, Biolegend, 5 μg/ml). Following incubation, cells were fixed and permeabilized with CytoFix/CytoPerm (BD Biosciences) at room temperature for 20 min. Cells were then washed three times with 100 μl PBS 1% BSA 0.1% Saponin (Sigma). Immunostaining for actin was performed by incubating with phalloidin Alexa Fluor 568 (Invitrogen) at 1:100 in the dark for 1 hr at room temp. Coverslips were mounted to slides using Prolong Glass antifade reagent (ThermoFisher Scientific) or Vectashield (Vector Laboratories). Images were acquired on a GE Deltavision OMX SR equipped with a 60X 1.42 NA APO objective and 3 PCO Chip sCMOS cameras. 3D images were captured with 0.125 μm steps with a pixel size of 0.079 μm. SIM images were reconstructed with GE SoftWoRX software using 3 orientations and 5 phase shifts with a Wiener filter constant of 0.005 and negative values not discarded.

### Image Analysis

Fiji (34) was used to process and analyze all confocal and structured illumination microscopy images. Line profiles were drawn using the ‘Line Profile’ tool and the MFI of each channel was exported to Prism (GraphPad Software) for normalized MFI comparison of CD56 and actin over the length of the line. For measurement of CD56 accumulation at the immune synapse in target cell conjugates, intensity measurements (AU) were made using the area of a region of interest (μm^2^) multiplied by the mean fluorescence intensity after auto thresholding. Granule convergence measurements were made using the length of a line drawn from the centroid of perforin to the MTOC while MTOC polarization measurements used a line drawn from the MTOC to the center of the immune synapse. Granule polarization measurements were taken by making a 3D projection of a cell on activated glass and dividing the image into one half defined as the cell and the other as the synapse (closest to the plane of the glass). MFI of perforin was measured and plotted for comparison between sections of the cell and between cell types.

### Statistical Analyses

Prism 8.0 (GraphPad software) was used for statistical analyses. Student’s t-tests with Welch’s correction or one-way ANOVAs with multiple corrections were used for analysis throughout the study, unless otherwise stated. Measurement differences that produced a p-value of <0.5 were deemed statistically significant. To generate technical repeats, all experiments using cell lines were performed on different days with different passages of cells. In some cases, data are shown as SuperPlots as described by Lord et al. (35).

## ACKNOWLEDGEMENTS

These studies used the resources of the Herbert Irving Comprehensive Cancer Center Flow Cytometry Shared Resources funded in part through Center Grant P30CA013696. This work was supported in part by NIH-NIAID R01AI137073 to EMM.

